# Prevalence and genotype-specific distribution of Human papillomavirus in Burundi according to HIV status and urban or rural residence and its implications for control

**DOI:** 10.1101/488429

**Authors:** Zacharie Ndizeye, Davy Vanden Broeck, Ramokone Lebelo Lisbeth, Johannes Bogers, Ina Benoy, Jean-Pierre Van Geertruyden

## Abstract

**Background:** Human papillomaviruses are the most important causative agents for invasive cervical cancer development. HPV type-specific vaccination and HPV cervical cancer screening methods are being widely recommended to control the disease but the epidemiology of the circulating HPV types may vary locally. The circulating HPV-strains have never been assessed in Burundi. This study determined the prevalence and genotype-specific distribution of HPV in four different strata in Burundi: HIV-infected or non-infected and women living in rural or urban areas. Implications for HPV diagnosis and vaccine implementation was discussed.

**Methods:** Four cross-sectional surveys were conducted in Burundi (2013 in a rural area and 2016 in urban area) among rural and urban HIV-infected and uninfected women. Cytology and HPV genotyping was performed to screen women for cervical cancer lesions. Risk factors for HPV infection and cervical cancer lesions were determined using logistic regression model.

**Results:** HPV prevalence was very high in urban area with significant differences between HIV-positive and negative women (p<0.0001). In fact, 45.7% of HIV-positive participants were infected with any HPV type and all were infected with at least one HR/pHR-HPV type. Among the HIV-negative participants, 13.4% were HPV-infected, of whom, only 4 women (2.7%) were infected with HR/pHR-HPV types. In rural, HPV infection did not significantly differ between HIV-positive and negative women (30.0% and 31.3% respectively; p=0.80).

In urban, multiple infections with HR/pHR-HPV types were detected in 13.9% and 2.7% among HIV-positive and negative women respectively (p<0.0001), whereas in rural, multiple infections with HR/pHR-HPV types were detected in 4.7% and 3.3% of HIV positive and negative women respectively (p=0.56).

The most prevalent HR/pHR-HPV types in HIV-positive urban women were HPV 52, 51 and 56. In the HIV-negative urban women, the most prevalent HR/pHR-HPV types were HPV 66, 67 and 18. In HIV-positive rural women, the most prevalent HR/pHR-HPV types were HPV 66, 16 and 18. In the HIV-negative rural women, the most prevalent HR/pHR-HPV types were HPV 16, 66 and 18. Independent risk factors associated with cervical lesions were HPV and HIV infections.

**Conclusions:** There is a high burden of HR/pHR-HPV infections, in particular among HIV-infected urban women. The study points out the need to introduce a comprehensive cervical cancer control program adapted to the context. This study shows that the nonavalent vaccine covers most of the HR/pHR-HPV infections in rural and urban areas among HIV-infected and uninfected women.

## Background

Since about three decades, human papillomaviruses (HPVs) have been firmly proven to be the most important etiologic agents for the development of invasive cervical cancer (ICC) [1-3]. Worldwide, HPVs are known to be one of the most common Sexually Transmitted Infections (STIs) and HPV prevalence peaks soon after sexual debut during adolescence and decreases thereafter with increasing age [4;5].

HPVs are small double-stranded DNA viruses, with a large epithelial tropism. Basal epithelial cells are infected with HPVs, causing benign and malignant lesions of the skin and the ano-genital mucosae and the upper aero-digestive tract [6]. Studies on HPV epidemiology and their mechanistic evidence have led to their classification into three groups: (1) HPV types 16, 18, 31, 33, 35, 39, 45, 51, 52, 56, 58 and 59 are known as carcinogenic, also named high-risk (HR)-HPV types; (2) HPV types 26, 53, 66, 67, 68, 70, 73 and 82, are classified as probably/possibly high-risk carcinogenic (pHR)-HPV types and (3) other HPV types such as HPV 6 and 11 are classified as low-risk (LR)-HPV types [7]. However, there is a growing literature providing evidence that the classified (pHR)-risk HPV types may also have to be considered as (HR)-HPV types [7-9].

Risk factors for HPV infection have been documented and include infection with other STIs (including HIV), high number of lifetime sexual partners, early sexual debut and host susceptibility [6;10]. Non-sexual transmission routes have also been documented but account for a small minority of HPV infections. They include perinatal transmission and, possibly, transmission by medical procedures and fomites [6].

Both HIV and HPV are STIs and infection with either one of the two viruses may facilitate transmission of the other [14;15]. Furthermore, HPV infections are more persistent in HIV-infected women compared to HIV-negative women, and as a consequence, cervical lesions are more frequent in HIV-positive women than in HIV-negative women [16;17]. Thus, HIV-infected women are at higher risk of developing ICC compared to HIV-uninfected women [11-13]. Prevention of ICC in sub-Saharan Africa (SSA) through screening with the conventional Pap smear method or other alternative techniques such as visual inspection with Acetic Acid/Lugol’s iodine is challenging and barely rolled out on public health scale [18;19]. New control strategies such as HPV screening with rapid molecular HPV tests and HPV vaccination are emerging and may be promising complementary tools for the Low and Middle Income Countries (LMICs) [20].

Currently, there are three prophylactic HPV vaccines licensed: (1) Cervarix^®^ (GlaxoSmithKline, Brentford, UK); a bivalent vaccine targeting HPV16/18; (2) Gardasil^®^ (Merck Inc, NY, USA), a quadrivalent vaccine targeting HPV6/11/16/18 and (3) Gardasil-9^®^ (Merck Inc., NY, USA), a nonavalent vaccine targeting HPV 6/11/16/18/31/33/45/52/58 [21].

In Burundi, ICC represents the most common female cancer, accounting for approximately 39% of all female cancers [22]. It is responsible, each year, for an estimated 1 421 new cases and 1 080 deaths in Burundi, representing an annually age-standardised incidence and mortality rates of 49.3 and 39.3/100,000 women respectively [22]. Burundi does not have any cervical cancer screening programme because of a variety of factors including lack of adequate infrastructure, insufficiently qualified staff and insufficient investment in resources for pap-smears, biopsies and colposcopy [23;24].

Since end of 2016, a demonstration project on HPV vaccination started which was limited to two districts where girls aged 9–13 years old are being vaccinated with Cervarix^®^. It is expected to expand at national level in a second phase after evaluation of this demonstration project [25]. However, the impact of HPV vaccination will be only fully realized several decades after a vaccination programme is instituted. Further, immunization can be ineffective due to insufficient coverage, missed follow-up doses, strain incompatibility and cost. Indeed, it remains challenging to guarantee sufficient coverage and ensure all girls of appropriate age to be vaccinated. Apart from report on cross-protection [26], the bivalent vaccine chosen by Burundi offers protection against the two most prevalent HR-HPV types 16 and 18. Both genotypes are expected to be responsible for 70% of all cancer cases, implying that at least 30% of all cervical cancer cases might not be covered by this prophylactic vaccination.

To date, information about the epidemiology of the circulating HPV strains in Burundi has not yet been ever documented. In the current context of type-specific HPV vaccination and of HPV-based cervical cancer screening, information about the prevalence and the circulating HPV types and their relative contribution to ICC is of great importance to assist in planning for vaccine, cervical cancer screening implementation as well as to monitor the potential impact on circulating HPV types after vaccination.

Hence, this study aimed to document the prevalence and genotype-specific distribution of cervical HPV types in both HIV-infected and uninfected Burundian women living in urban (Bujumbura) or in rural setting (Kirundo).

## Materials and methods

### Study design, setting and population

Four cross-sectional surveys were conducted: two were conducted from May to July 2013 in Kirundo, a rural health district in the northern part of Burundi. Another two were conducted from March to May 2016 in Bujumbura, the capital city of Burundi.

In the rural area, participants were women attending Kirundo District Hospital and ANSS (Association Nationale de Soutien aux Séropositifs et malades du SIDA)-Kirundo antenna. The province of Kirundo is divided into 4 health district zones but has only 2 district hospitals (Kirundo and Mukenke). Kirundo district hospital is located at the provincial town of Kirundo and is the biggest hospital in the province. Patients are referred from health centres in Kirundo district and the neighbouring Busoni and Vumbi health districts. Patients can also come directly to the hospital to consult a General Practitioner (GP). ANSS-Kirundo is a local NGO, champion in HIV-care in Kirundo and has the longest active list of HIV-patients coming from the 4 health districts and few from the neighbouring Ngozi province. It is located near Kirundo district hospital.

In the urban area of Bujumbura, participants were women attending an HIV clinic, located in the University Teaching Hospital of Bujumbura, which follows up around 3500 HIV-positive patients. Participants were also women attending a reputed family planning centre, ABUBEF (Association Burundaise pour le Bien-Etre Familial), in Bujumbura with clients from all neighbourhoods and all social classes.

The study was approved by the Burundian National Ethics Committee. A written informed consent to participate in the survey (translated in the local language) was obtained and signed by all participants before being enrolled in the study. Information collected was kept confidential by the use of codes.

### Recruitment procedures

General information about the study objectives were given to all participants every morning in the waiting room. During the consultation in the outpatient department, a GP gave clear and detailed information on the study objective and procedures, and proposed the women to participate in the study. Women who signed an informed consent were assisted by the GP to complete a short risk factor questionnaire and blood was collected for HIV testing. Women who declared being HIV-negative or with an unknown HIV status received a pre-test counselling before HIV-testing. A post-test counselling was done before giving back the result. Those who tested HIV-positive were referred to any HIV clinic of their choice for follow up. Women known to be HIV-positive, followed up at the University HIV clinic or ANSS were not retested for HIV.

After filling the questionnaire, a gynaecologic examination was conducted. Cervical samples were collected using a cytobrush^®^ (Cervex-Brush combi, Rovers Medical Devices B.V., The Netherlands) and placed into liquid-based cytology (LBC) medium (ThinPrep, Hologic). Vials were stored at room temperature until shipment. Specimens were sent to the Department of Virology at Sefako Makgatho Health Science University (SMU) in Pretoria, South Africa and to the Laboratory of Molecular Pathology, AML, Sonic Healthcare, Antwerp, Belgium for HPV genotyping and cytology reading.

### Inclusion and exclusion criteria

Any woman (or girl) aged between 17-65 years, declaring having had vaginal sexual intercourse and agreeing to participate could be included in the study. Exclusion criteria were pregnancy, menstrual period, vaginal discharge and hysterectomy.

### Sample size determination

We estimated HPV prevalence in HIV-negative and HIV-positive women to be 21% and 36.3% respectively [16;22;27]. With a power of 80% for a confidence level of 95%, the minimum sample size required was 149 subjects per stratum. The calculation was done using the EpiInfo7 software. We therefore decided to investigate 150 HIV positive and 150 HIV negative women in both the rural and urban settings.

### HPV testing and liquid based cytology processing

We initially planned that all samples would be tested at SMU, South Africa. At the time of sample collection of urban specimens, we realized that the laboratory was overloaded and therefore have decided to send urban samples at AML, Antwerp in Belgium.

For the 300 samples from the urban area, thin-layer slides were prepared with the ThinPrep^®^ 5000 Processor with Autoloader System (Hologic Inc, Marlborough, US) and stained with the Papanicolaou stain on the Tissue-Tek^®^ Prisma and Film Automated Slide Stainer and Coverslipper (Sakura Finetek Europe B.V., Netherlands). After scanning of the slides with the ThinPrep^®^ Imaging System, cytology reading was performed by image-guided screening, with prior knowledge of HPV infection status. Cytological diagnoses were reported according to the Bethesda 2001 terminology system as 1) normal, 2) atypical squamous cells of undetermined significance; atypical squamous cells cannot exclude high-grade lesion; atypical glandular cells; low-grade squamous intraepithelial lesions (ASCUS/ASC-H/AGC/LSIL), 3) high-grade squamous intraepithelial lesions (HSIL), or 4) invasive cancer. After the LBC preparations are made, 800μL of the remaining cell suspension was used for DNA extraction. Cytology reading was performed only on the 300 urban samples due to budget constraints.

### HPV type-specific detection

For the urban samples, DNA isolation from liquid-based cytology was performed on the Medium Throughput Automation (MTA) (Hologic Inc) with the Genfind® DNA extraction kit. Subsequently, the DNA is amplified using a series of real-time qPCR reactions in the LightCycler 480 (Roche) as previously described by Micalessi et al [28]. Briefly, the RIATOL qPCR HPV genotyping assay is a clinically validated, laboratory developed test, which amplifies 18 HPV types: HPV 6E6, 11E6, 16E7, 18E7, 31E6, 33E6, 35E6, 39E7, 45E7, 51E7, 52E7, 53E6, 56E7, 58E7, 59E7, 66E6, 67 L1 and 68E7. Real-time quantitative PCR for β-globin was always performed and was used as a proxy for the quality of sampling.

The 300 specimens from rural area were tested at the Department of Virology, SMU, Pretoria, RSA. HPV DNA was extracted using AmpliLute liquid media Extraction kit following manufacturer’s instructions (Roche Molecular Systems, California, USA). HPV DNA was detected following conventional nested HPV. PCR was performed with MY09/MY11 (450 bp) and GP5+/GP6+ (150 bp). First round PCR was carried out in a final reaction of 50 μl containing 1 X buffer II (Bioline, Luckenwalde, Germany), 1.5 mM MgCl2 (Bioline, Luckenwalde, Germany), 200 µM of each deoxynucleoside triphosphate (dNTP), 0.2 µM of MY09/MY11 primers and 0.2 µM PCO3 and PCO4 primers (110 bp) (Inqaba Biotechnological Industry, Pretoria, South Africa), 1 unit of Bio Taq DNA polymerase (Bioline, Hilden, Germany), and 10 μl of DNA template. The cycling conditions included an initial denaturation step for 2 minutes at 94 °C followed by 40 cycles at 94 °C for denaturation, 30 seconds at 55 °C for annealing, 1 minute at 72 °C for elongation and a final elongation for 5 minutes at 72°C. The PCR product detected using gel electrophoresis (2% w/v agarose in Tris-acetate EDTA) and visualised using ethidium bromide. Bands of the appropriate size were identified by comparison with a DNA molecular weight marker and the gel was viewed using a Gel DOC system (Syngene, Europe). HPV genotyping was performed using Linear array genotyping test that identifies 37 HPV genotypes (6, 11, 16, 18, 26, 31, 33, 35, 39, 40, 42, 45, 51, 52, 53, 54, 55, 56,58, 59, 61, 62, 64, 66, 67, 68, 69, 70,71, 72 73, 81, 82, 83, 84, 89, IS39) as previously described by Coutlee et al. [29].

Briefly, 50μl of amplicon was added to 50μl of a working master mix containing MgCl2, Amplitaq® GOLD DNA polymerase, Uracil-N-glycosilase, deoxynucleotides, PGMY and β globin primers. The PCR amplification was performed using the gold-plated 96-well Gene Amp PCR System 9700 (Applied Biosystems, Foster City, California, USA) according to manufacturer instructions. Positive reactions appeared as blue lines and were interpreted using the LA HPV GA reference guide.

### Statistical analysis

Proportions were compared using Pearson’s Chi-square test (or Fisher’s exact test when appropriate). Ttest was used to compare means of normally distributed variables (or Kruskal-Wallis test when appropriate). Bivariate analysis was performed to generate odds ratios (ORs) with their 95% confidence interval (CI) to analyse the relationship between each sociodemographic variable and cervical cancer lesions or with HPV infection status. Variables with a bivariate p-value <0.20 were entered in a multivariate logistic regression model to determine adjusted odds ratios (AORs) with their 95% CIs. Analysis was conducted using EpiInfo 3.5.4 software and α-error margin of 5% was considered significant.

## RESULTS

### Participants and their socio-demographic characteristics

In rural area, we enrolled 150 HIV-negative and 150 HIV-positive women. The mean age of the participants was 39.9 years (SD=8.3) and 36.4 years (SD=7.9) for HIV-positive and HIV-negative women respectively (p=0.0002). Profession categories and marital status differed between HIV-positive and negative women (p<0.0001 for both). Farmers represented 71.3% and 50% in HIV-positive and negative women respectively. The other HIV-negative women were ‘’employees/shopkeepers’’ (47.3%) or ‘’housewife/Non-working/student/other’’ (2.7%). A hundred thirty-one HIV-positive women (87.3%) was married compared to seventy-eight (52%) in the HIV-positive group. In the HIV-positive group, 38% were “widowed or divorced” compared to 10% in HIV-negatives. HIV-positive women started earlier sexual intercourse, got married earlier, got pregnant earlier and had a higher median number of sexual partners compared to their HIV-negatives counterparts (all p<0.001) (Table 1).

In urban, a total of 151 HIV-positive and 149 HIV-negative participants were enrolled. The mean age of our participants was 41.1 years (SD=9.7) and 39.7 years (SD=8.7) for HIV-positive and HIV-negative women (p=0.19). Profession categories and marital status differed between HIV-positive and negative women (p<0.0001 for both). In the HIV-positive group, majority (42.4%) was in the category ‘’housewife/Non-working/student/other’’. In the HIV-negative group, 77.2% of participants were employees (in public or private) or shopkeepers. In the HIV-negative group, 91.9% were married versus 53% in the HIV-positive group. A high proportion of HIV-positive women were widowed/divorced (36.4%) whereas only 4% who belonged to this category were in the HIV-negative group. HIV-positive women started earlier sexual intercourse, got married earlier, got pregnant earlier and had a higher median number of sexual partners compared to their HIV-negative counterparts (all p<0.001) (Table 1).

**Table 1:**
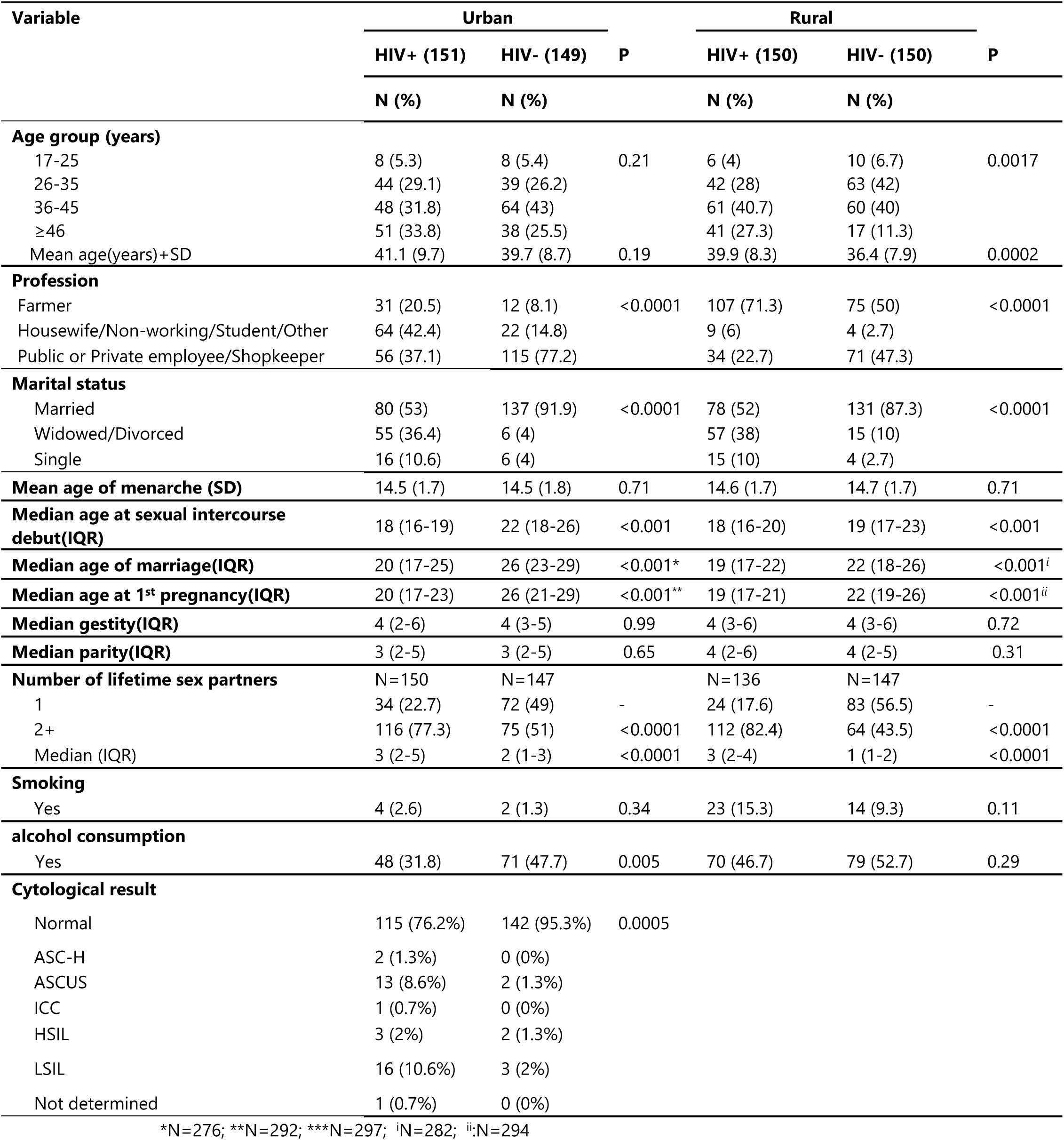
Baseline socio-demographic characteristics of the participants by study area and HIV status, among 600 women, Burundi, 2016.

### HPV prevalence

In rural, all participants had valid data on HPV genotyping. The HPV prevalence (any) was 30% (45/150) and 31.3% (47/150) among the HIV-positive and negative women respectively (p=0.80). Multiple HPV infection was 11.3% and 14% among the HIV-positive and negative women respectively (p=0.49). The HR/pHR-HPV prevalence was 18.7% and 17.3% in HIV-positive and negative respectively (p=0.76). Multiple HR/pHR-HPV infections was 4.7% versus 3.3% among HIV-positive and negative women respectively (p=0.56).

In the HIV-positive group, infection with any HR-HPV, pHR-HPV and LR-HPV type was 14.7%, 7.3% and 19.3% respectively. In HIV-negative women, infection with any HR-HPV, pHR-HPV and LR-HPV type was 12%, 6.7% and 24.7% respectively (Table 2). Infection with HPV 16 or 18 was 8% versus 6.7% among HIV-positive and negative women respectively (p=0.50). The prevalence of the HR-HPV types included in the Gardasil-9 vaccine was 14% versus 11.3% among HIV-positive and negative women respectively (p=0.49). The most frequent HR-HPV types in the HIV-positive group were HPV 16 (4%), 18 (4%), followed by HPV 33 (3.3%) and HPV 58 (2%). The most frequent pHR-HPV types were HPV 66 (4.7%) and HPV 70 (3.3%). Among the LR-HPV types, the most frequently isolated were HPV 11 (15.3%) and HPV 6 (2%).

**Table 2:** Prevalence of human papillomavirus by HIV status in rural and urban areas among 600 women in Burundi.

| Type de HPV | Rural |  |  | Urban |  |  |  |  |  |  |  | P<sup>a</sup> |
| --- | --- | --- | --- | --- | --- | --- | --- | --- | --- | --- | --- | --- |
|  | HIV positive<br>(N=150) | HIV negative<br>(N=150) | P | HIV positive women |  |  |  | HIV negative women |  |  |  |  |
|  |  |  |  | Normal<br>(N=115) | ASCUS/LSIL<br>(N=29) | HSIL/ASC-H/<br>ICC (N=6) | Total<br>(N=151) | Normal<br>(N=142) | ASCUS/LSI<br>L (N=5) | HSIL/ASC-H/<br>ICC (N=2) | Total<br>(N=149) |  |
|  |  |  |  | N (%) | N (%) | N (%) | N (%) | N (%) | N (%) | N (%) | N (%) |  |
| Positive | 45 (30.0) | 47 (31.3) | 0.80 | 33 (28.7) | 29 (100) | 6 (100) | 69* (45.7) | 13 (9.2) | 5 (100) | 2 (100) | 20 (13.4) | <0.0001 |
| Multiple | 17 (11.3) | 21 (14.0) | 0.49 | 8 (7) | 11 (37.9) | 2 (33.3) | 22 (14.6) | 2 (1.4) | 2 (40) | 0 (0) | 4 (2.7) | <0.0001 |
| HR/pHR-HPV (any) | 28 (18.7) | 26 (17.3) | 0.76 | 33 (28.7) | 29 (100) | 6 (100) | 69* (45.7) | 12 (8.4) | 5 (100) | 2 (100) | 19 (12.8) | <0.0001 |
| Multiple HR/pHR-HPV | 7 (4.7) | 5 (3.3) | 0.56 | 8 (7) | 10 (34.5) | 2 (33.3) | 21* (13.9) | 2 (1.4) | 2 (40) | 0 (0) | 4 (2.7) | <0.0001 |
| HPV 16-18 | 12 (8.0) | 9 (6.0) | 0.50 | 6 (5.2) | 3 (10.3) | 4 (66.7) | 14* (9.3) | 1 (0.7) | 1 (20) | 2 (100) | 4 (2.7) | 0.016 |
| HPV 16-18-31-33-45-52-58 | 21 (14.0) | 17 (11.3) | 0.49 | 18 (15.7) | 14 (48.3) | 5 (83.3) | 38* (25.2) | 3 (2.1) | 4 (80) | 2 (100) | 9 (6) | <0.0001 |
| 35-39-51-53-56-59-66-67-68 | 8 (5.3) | 11 (7.3) | 0.50 | 18 (15.7) | 22 (75.9) | 2 (33.3) | 43* (28.5) | 9 (6.3) | 2 (40) | 0 (0) | 11 (7.4) | <0.0001 |
| High risk |  |  |  |  |  |  |  |  |  |  |  |  |
| 16 | 6 (4.0) | 7 (4.7) |  | 2( 1.7) | 1 (3.4) | 3 (50) | 7 (4.6) | 0 (0) | 0 (0) | 1 (50) | 1 (0.7) |  |
| 18 | 6 (4.0) | 3 (2.0) |  | 4 (3.5) | 3 (10.3) | 1 (16.7) | 8 (5.3) | 1 (0.7) | 1 (20) | 1 (50) | 3 (2) |  |
| 31 | 2 (1.3) | 1 (0.7) |  | 3 (2.6) | 1 (3.4) | 1 (16.7) | 6 (4) | 0 (0) | 1 (20) | 0 (0) | 1 (0.7) |  |
| 33 | 5 (3.3) | 1 (0.7) |  | 1 (0.9) | 0 (0) | 0 (0) | 1 (0.7) | 0 (0) | 0 (0) | 0 (0) | 0 (0) |  |
| 35 | 0 (0.0) | 2 (1.3) |  | 2 (1.7) | 3 (10.3) | 1 (16.7) | 6 (4) | 0 (0) | 0 (0) | 0 (0) | 0 (0) |  |
| 39 | 0 (0.0) | 0 (0.0) |  | 2 (1.7) | 1 (3.4) | 0 (0) | 3 (2) | 2 (1.4) | 0 (0) | 0 (0) | 2 (1.3) |  |
| 45 | 0 (0.0) | 2 (1.3) |  | 2 (1.7) | 1 (3.4) | 0 (0) | 4 (2.6) | 2 (1.4) | 1 (20) | 0 (0) | 3 (2) |  |
| 51 | 1 (0.7) | 0 (0.0) |  | 4 (3.5) | 6 (20.7) | 0 (0) | 12 (7.9) | 1 (0.7) | 0 (0) | 0 (0) | 1 (0.7) |  |
| 52 | 0 (0.0) | 2 (1.3) |  | 7 (6.1) | 6 (20.7) | 0 (0) | 13 (8.6) | 0 (0) | 0 (0) | 0 (0) | 0 (0) |  |
| 56 | 1 (0.7) | 0 (0.0) |  | 3 (2.6) | 7 (24.1) | 0 (0) | 10 (6.6) | 0 (0) | 0 (0) | 0 (0) | 0 (0) |  |
| 58 | 3 (2.0) | 2 (1.3) |  | 4 (3.5) | 4 (13.8) | 1 (16.7) | 9 (6) | 0 (0) | 2 (40) | 0 (0) | 2 (1.3) |  |
| 59 | 0 (0.0) | 0 (0.0) |  | 2 (1.7) | 0 (0) | 0 (0) | 2 (1.3) | 1 (0.7) | 0 (0) | 0 (0) | 1 (0.7) |  |
| Any HR | 22 (14.7) | 18 (12.0) |  | 28 (24.3) | 26 (89.6) | 6 (100) | 61* (40.4) | 5 (3.5) | 4 (80) | 2 (100) | 11 (7.4) |  |
| Possibly high risk |  |  |  |  |  |  |  |  |  |  |  |  |
| 26 | 1 (0.7) | 1 (0.7) |  |  |  |  |  |  |  |  |  |  |
| 53 | 0 (0.0) | 1 (0.7) |  | 1 (0.9) | 4 (13.8) | 0 (0) | 5 (3.3) | 2 (1.4) | 0 (0) | 0 (0) | 2 (1.3) |  |
| 66 | 7 (4.7) | 6 (4.0) |  | 1 (0.9) | 2 (6.9) | 0 (0) | 4 (2.6) | 2 (1.4) | 2 (40) | 0 (0) | 4 (2.7) |  |
| 67 | 1 (0.7) | 2 (1.3) |  | 3 (2.6) | 1 (3.4) | 0 (0) | 4 (2.6) | 4 (2.8) | 0 (0) | 0 (0) | 4 (2.7) |  |
| 68 | 0 (0.0) | 0 (0.0) |  | 2 (1.7) | 3 (10.3) | 0 (0) | 6 (4) | 0 (0) | 0 (0) | 0 (0) | 0 (0) |  |
| 70 | 5 (3.3) | 1 (0.7) |  |  |  |  |  |  |  |  |  |  |
| Any pHR | 11 (7.3) | 10 (6.7) |  | 6 (5.2) | 8 (27.6) | 0 (0%) | 15* (9.9) | 7 (4.9) | 2 (40) | 0 (0) | 9 (6.0) |  |
| Low risk |  |  |  |  |  |  |  |  |  |  |  |  |
| 6 | 3 (2.0) | 6 (4.0) |  | 0 (0) | 0 (0) | 0 (0) | 0 (0) | 1 (0.7) | 1 (20) | 0 (0) | 2 (1.3) |  |
| 11 | 23 (15.3) | 30 (20.0) |  | 0 (0) | 1 (3.4) | 0 (0) | 1 (0.7) | 0 (0) | 0 (0) | 0 (0) | 0 (0) |  |
| 40 | 2 (1.3) | 1 (0.7) |  |  |  |  |  |  |  |  |  |  |
| 54 | 1 (0.7) | 3 (2.0) |  |  |  |  |  |  |  |  |  |  |
| 55 | 1 (0.7) | 1 (0.7) |  |  |  |  |  |  |  |  |  |  |
| 81 | 2 (1.3) | 0 (0.0) |  |  |  |  |  |  |  |  |  |  |
| 83 | 0 (0.0) | 1 (0.7) |  |  |  |  |  |  |  |  |  |  |
| 84 | 1 (0.7) | 0 (0.0) |  |  |  |  |  |  |  |  |  |  |
| Any LR | 29 (19.3) | 37 (24.7) |  | 0 (0) | 1 (3.4) | 0 (0) | 1 (0.7) | 1 (0.7) | 1 (20) | 0 (0) | 2 (1.3) |  |

\*Including 1 participant with undetermined cytological result. <sup>a</sup> comparison of total HIV+ and total HIV- in urban

In the HIV-negative group, the most frequent HR-HPV types isolated were HPV 16 (4.7%), followed by HPV 18 (2%). The most frequent pHR-HPV types were HPV 66 (4%) and HPV 67 (1.3%). The most frequent LR-HPV types were HPV 11 (20%) and HPV 6 (4%).

Table 2 also presents HPV prevalence results for the urban area. The overall HPV prevalence (any HPV) was 45.7% and 13.4% among HIV-positive and HIV-negative women (p<0.0001). HPV prevalence is presented by cytological results and by HIV-status. Multiple infection with any HPV type was 14.6% and 2.7% among HIV-positive and negative women respectively (p<0.0001). The overall prevalence of HR/pHR-HPV infection was 45.7% and 12.8% with a multiple infection of 13.9% and 2.7% among HIV-positive and negative women respectively (p<0.0001 for both variables). In the HIV-positive group, HR/pHR-HPV prevalence was 28.7% (33/115), 100% (29/29) and 100% (6/6) among women with normal cytology, ASCUS/LSIL and HSIL/ICC/ASC-H respectively.

In the HIV-negative group, HR/pHR-HPV prevalence was 8.4% (12/142), 100% (5/5) and 100% (2/2) in women with normal cytology, ASCUS/LSIL and HSIL/ASC-H/ICC respectively.

Most prevalent HR/pHR-HPV types, in HIV-positive women, were HPV 52, 18, 51 and 58 among women with normal cytology; HPV 56, 51 and 52 in women with ASCUS/LSIL and HPV16 and 18 in women with HSIL/ASC H/ICC. In the HIV-negative women, the most prevalent HR/pHR-HPV types were HPV 67, 39, 45 and 66 in women with normal cytology; HPV 58 and 66 in women with ASCUS/LSIL and HPV 16 and 18 in women with HSIL/ASC-H/ICC.

Figure 1 (part A) presents the age-specific prevalence of HR/pHR-HPV by HIV status and by study area. It appears that HIV-positive urban women are highly infected with HR/pHR-HPV types than their counterparts in rural across all age groups. Rural HIV-negative women have slightly higher prevalence compared to their counterparts in urban in all age groups, except in the younger women who were not infected with any HR/pHR-HPV type.

**Figure 1:**
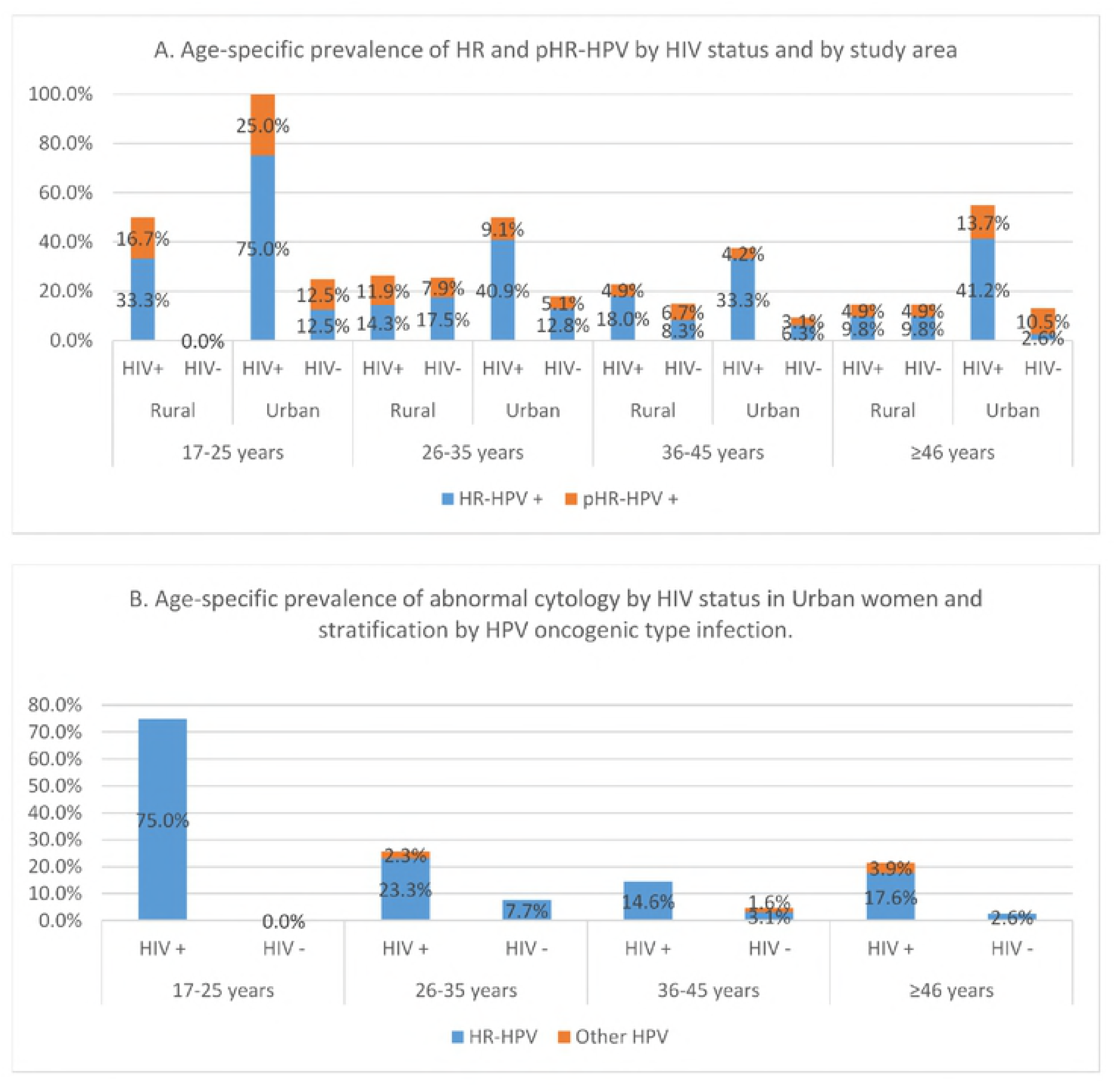
Age-specific prevalence of HPV (A) and of abnormal cytological results (B) stratified by HIV status in rural and urban women, Burundi.

Overall, among these HPV infections, HR-HPV types are the most predominant. In urban, HR/pHR-HPV infections decreased with increasing age, with a second peak in the age group ≥46 years, both in HIV-positive and negative women. In rural, the decrease in HR/pHR-HPV prevalence was slow compared to urban across all age groups.

Figure 1 (part) B presents the age-specific prevalence of abnormal cytological results in urban area. Further, it also presents the HR/pHR-HPV infections among these women with abnormal cytological results and it clearly appears that HR-HPV infections are the most predominant among women with abnormal cytological results. Cytological abnormalities also seem to decrease with increasing age, with a second peak in age group ≥46 years as it was also for HPV prevalence.

Cytology results were available for all urban women except one HIV-positive woman with an undetermined cytological result. Cytological abnormalities were more frequent in HIV-positive women compared to HIV-negative women (p=0.0005).

In the HIV-positive group, among 150 women with a valid result on LBC, 23.3% (35/150) had an abnormal cytological result, including 8.7% (13/150) with ASCUS, 10.6% (16/150) with LSIL, 1.3% (2 patients) with ASC-H, 2% with HSIL and 1 patient with ICC. In the HIV-negative group, among 149 women with valid results on LBC, 4.7% (7/149) had abnormal cytological results including 2 patients (1.3%) with ASCUS, 3 patients (2%) with LSIL and 2 patients (1.3%) with HSIL.

In the bivariate analysis, abnormal cytology was significantly associated with HR/pHR-HPV infection (OR=199; 95% CI:26.7-1483), age (older women i.e. age groups 36-45 years and ≥46 years being protected compared to younger women aged 17-25 years), number of lifetime sexual partners (having had two or more lifetime sexual partners was associated with a higher risk of abnormal cytology, OR=2.57; 95% CI: 1.14-5.80 compared to having had only 1 sexual partner), HIV infection status (being HIV-infected was associated with 6 times higher risk of having abnormal cytology than those who are HIV-uninfected, OR=6.17 95% CI: 2.64-14.42) and profession (being an employee or a shopkeeper was a protective factor of having abnormal cytology compared to farmers, OR=0.30 95%CI: 0.13-0.69). After having adjusted for age, marital status, HIV infection, profession and number of lifetime sexual partners, HR/pHR-HPV infection remained the only stronger significant predictor for abnormal cytology (OR=162.54; 95%CI: 20.9-1261.4).

Table 4 presents the predictors of HPV infection in urban and rural. In urban, abnormal cytology and HIV infection were the most important contributor to a positive HPV diagnosis (OR=148.72; 95%CI: 19-1164.6 and 4.06 95%CI: 1.78-9.22 respectively), whereas in rural, there was no significant contributor identified, after having controlled for other covariates (age, lifetime number of sex partners, profession, parity and marital status).

Figure 2 presents the distribution of HR/pHR-HPV among HPV-positive participants by HIV status. In the HIV-positive rural women, infection with HPV 16 or 18 represented 32.1% of all HR/pHR-HPV infections. In the HIV-negative rural women, infection with HPV 16 or 18 represented 30.8% of all HR/pHR-HPV infections. Infections with HPV 16/18/31/33/45/52/58 represented 60.7% and 53.9% of the circulating HR/pHR-HPV infections among HIV-positive and negative rural women respectively.

**Figure 2:**
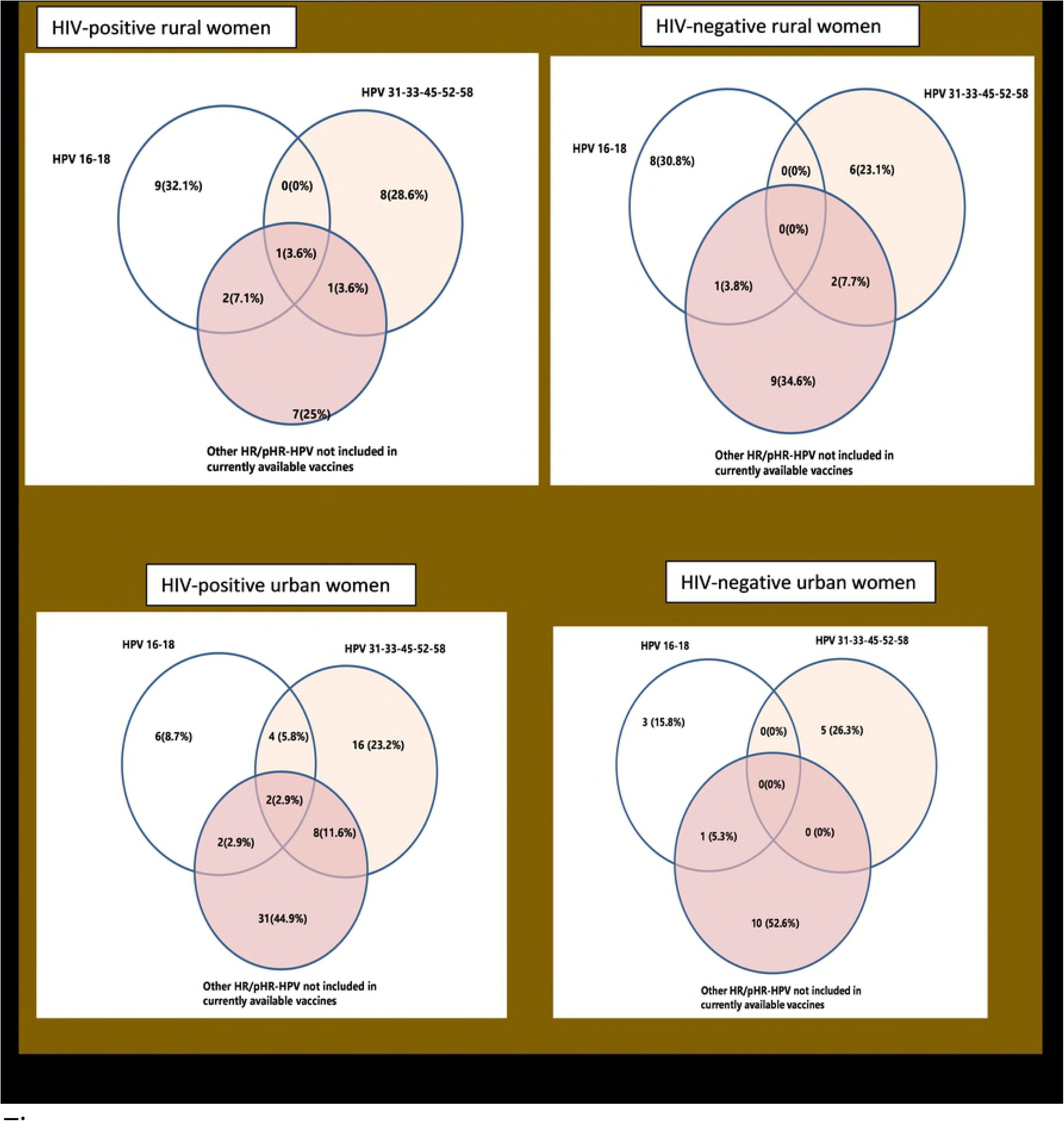
Distribution of HR/pHR-HPV types among HPV + women by study area and HIV status.

In urban women, HPV 16/18 infections represented 8.7% and 15.8% of all HR/pHR-HPV infections in HIV-positive and negative women respectively.

Infections with the HPV types included in the Nonavalent vaccine (i.e. HPV 16/18/31/33/45/52/58) represented 31.9% and 42.1% of all HR/pHR-HPV infections in HIV-positive and negative urban women respectively.

## Discussion

This study describes, to our knowledge, for the first time in Burundi the HPV prevalence and type-distribution according to HIV status in rural and urban areas. This may put in perspective diagnostic screening and vaccination program using type specific tools. Moreover, it presents HPV prevalence and type-distribution according to cytological findings in the urban area. These data are of young adult women, never screened for cervical lesions and before any national screening and/or vaccination programme has been rolled out.

According to Bruni L. et al. based on an extensive meta-analysis, the age distribution of cervical HPV infection in Africa is a bimodal curve, with a first peak at younger ages just after sexual debut, a lower prevalence plateau at middle ages, and a variable rebound at older ages(≥ 45 years) [30;31]. Our results from rural and urban follow a similar pattern, show high HPV prevalence in younger women (less than 35 years) with a relatively slow decrease with increasing age. The small differences with the pattern described by Bruni L. et al. and in other studies [32;33] could be due to small numbers in our analysis. Differences in age-specific HPV prevalence has also been reported in a study done in Kenya where HPV prevalence was similarly high across all age groups [34]. The authors hypothesised that the sexual behaviour of these women and/or their partners might not change with age to the same extent as in Western countries.

Our results show a high overall HPV prevalence ranging from of 13.4% to 45.7% in urban HIV-negative and positive women respectively (p<0.0001). Our data show significant differences in HR/pHR-HPV prevalence between HIV-positive and negative women in urban (45.7% and 12.8% respectively, p<0.0001). This pattern has been described in other studies [4;16;33;36]. Studies done in Africa among HIV-positive women have reported a HR/pHR-HPV prevalence ranging from 27% to 52% [4;32;36;37]. Two studies done in Rwanda by Ngabo et al. and Mukanyangezi et al. found HR/pHR-HPV prevalence in HIV-negative women of 8.1% and 20%, which is in line with our results [36;37]. In Tanzania, Dartell et al. reported a similar HR-HPV prevalence of 17.2% among HIV-negative women [32].

The overall HPV prevalence was 30 to 31.3% in rural women with no difference according to HIV status (p=0.80). This high HPV prevalence is in line with other studies conducted in East Africa [30;31;35]. However, HR/pHR-HPV prevalence of rural HIV-positive and negative women were not significantly different (18.7% and 17.3% respectively, p=0.76). This was an unexpected result and our hypothesis is that, HPV infection being related to sexual behaviour, this would imply that these HIV-positive rural women (probably also their partners) changed their sexual behaviour because of the illness and counselling received from HIV clinic health workers where they are followed up for their HIV treatment and did not get re-infected with new HPVs. This argument is reinforced by the observed consistently higher HPV prevalence of urban HIV-positive women across all age groups and suggests differences in sexual behaviour among HIV-positive rural and urban women.

Moreover, a selection bias cannot be ruled out as the participants were not randomly selected; all women were enrolled from the health facilities and thus may not represent the rural population. Therefore, more information in robust studies is needed to fully understand the pattern in this area. In the literature, lack of significant differences in HPV prevalence among HIV-positive and negative women has also been reported in a study done in a rural community in Zimbabwe [38].

As we expected, stronger independent predictors for abnormal cytology results were a positive HR/pHR-HPV infection and HIV infection (table 3). Indeed, persistent HR-HPV infection has been shown to be a necessary cause for the development of cervical lesions and HIV-infected women being at higher risk of developing cervical lesions. With the WHO last recommendations to start HAART among HIV-infected persons irrespective to their immunity level, we can only speculate that HAART will lead to a more rapid clearance of HR/pHR-HPV types among HIV-positive women, and as a consequence a reduction in incidence of cytology and histology diagnosed cervical lesions, as this has been reported by Kelly et al. [39] in their recent meta-analysis.

**Table 3:** Predictors of abnormal cytological results (ASCUS +) in urban women.

| Variable | Crude OR (95%CI) | P | Adjusted OR (95%CI) | p |
| --- | --- | --- | --- | --- |
| **HR/pHR-HPV infection** |  |  |  |  |
| No | - |  |  |  |
| Yes | 199.1 (26.7-1482.9) | <0.0001 | 162.54 (20.94-1261.42) | <0.0001 |
| **Age group (years)** |  |  |  |  |
| 17-25 | - |  | - |  |
| 26-35 | 0.34 (0.11-1.10) | 0.07 | 0.29 (0.05-1.66) | 0.17 |
| 36-45 | 0.16 (0.05-0.54) | 0.003 | 0.24 (0.04-1.44) | 0.12 |
| ≥46 | 0.26 (0.08-0.85) | 0.03 | 0.19 (0.03-1.18) | 0.07 |
| **HIV infection** |  |  |  |  |
| Yes | 6.17 (2.64-14.42) | <0.0001 | 1.69 (0.54-5.29) | 0.37 |
| No | - |  | - |  |
| **Number of lifetime sexual partners (N=297)** |  |  |  |  |
| 1 | - |  | - |  |
| ≥2 | 2.57 (1.14-5.80) | 0.02 | 1.69 (0.59-4.89) | 0.33 |
| **Profession** |  |  |  |  |
| Farmer | - |  | - |  |
| Housewife/Non-working/Student/other | 0.42 (0.17-1.05) | 0.06 | 0.23 (0.06-0.93) | 0.04 |
| Public or Private employee/Shopkeeper | 0.30 (0.13-0.69) | 0.005 | 0.41 (0.11-1.50) | 0.18 |
| **Marital status** |  |  |  |  |
| Married | - |  | - |  |
| Widowed/Divorced | 1.83 (0.88-3.87) | 0.11 | 0.94 (0.31-2.88) | 0.92 |
| Single | 3.74 (0.88-15.83) | 0.07 | 1.52 (0.20-11.27) | 0.68 |
| **age sexual debut** |  |  |  |  |
| <17 | - |  |  |  |
| ≥17 | 0.66 (0.33-1.34) | 0.25 |  |  |
| **Age of marriage (N=276)** |  |  |  |  |
| <17 | - |  |  |  |
| ≥17 | 0.97 (0.27-3.44) | 0.96 |  |  |
| **age at first pregnancy (N=292)** |  |  |  |  |
| <17 | - |  |  |  |
| ≥17 | 0.60 (0.23-1.57) | 0.30 |  |  |
| **Gestity** |  |  |  |  |
| <6 | - |  |  |  |
| ≥6 | 1.78 (0.89-3.57) | 0.10 |  |  |
| **Parity** |  |  |  |  |
| <6 | - |  | - |  |
| ≥6 | 0.80 (0.30-2.18) | 0.67 | 0.66 (0.16-2.84) | 0.58 |
| **Alcohol consumption** |  |  |  |  |
| Yes | - |  |  |  |
| No | 1.36 (0.68-2.71) | 0.38 |  |  |

**Table 4:**
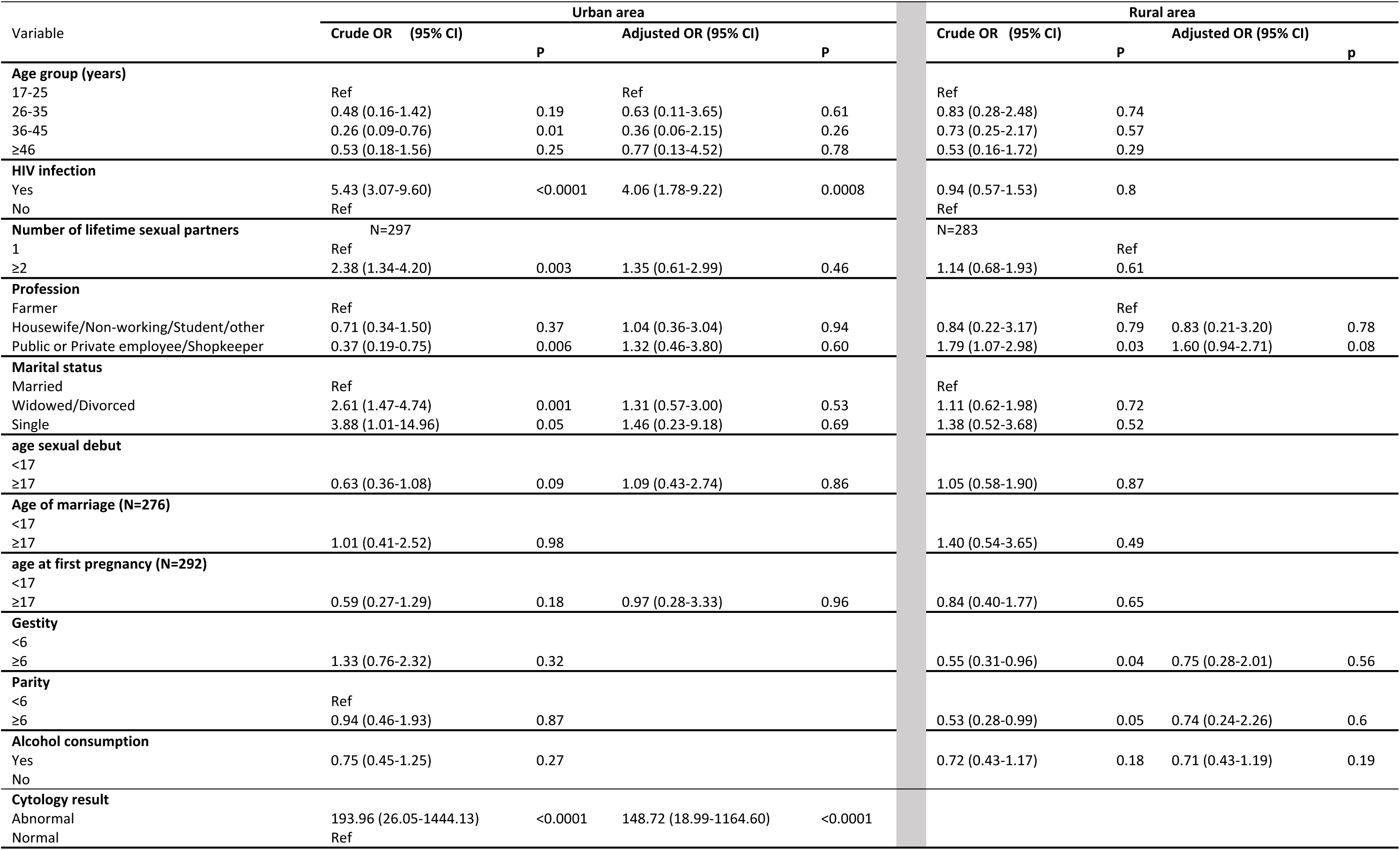
Predictors of the HPV (any) infection by study area (rural and urban) in women, Burundi.

Stratified by cytological results, our findings show a HPV prevalence of 28.7% and 9.2% in HIV-positive and negative women respectively with normal cytology, which is in line with data from East Africa showing HPV prevalence in women with normal cytology ranging from 3.2% to 41.4% [27;30;40;41].

HR/pHR-HPV prevalence in HIV-positive women with ASCUS/LSIL was 100% in our results and is higher than the prevalence found in a meta-analysis by Clifford M. et al [16]; which was 69.4%. We think that this difference could be due to large sample size in their study compared to ours. Another reason could be differences in the sensitivity of the HPV detection methods used in these studies as it has been reported [42]. Larger meta-analyses including studies on HR/pHR-HPV prevalence in the East-African region also showed similar results compared to ours among women with HSIL/ICC which ranged from 76.6 % to 96.6 %, and clearly showed an increase in HR/pHR-HPV prevalence as the severity of the cytological lesions increases [27;43].

### HPV strains and implications in the choice of HPV vaccines

Regarding the most isolated HPV strains among HIV-positive urban women with normal cytology, our findings are in agreement with results found in two robust meta-analyses presenting type-specific HPV prevalence in HIV-positive women with normal cytology in Africa despite some differences in ranking [16;41]. Consistent results with the literature are also found among women with LSIL/ASCUS. In women with high grade lesions (despite few numbers in our study, N=4), type-specific HPVs are concordant with results from other studies showing that the HPV 16 is by far the most prevalent HPV type [43;44].

In the HIV-negative group, this high frequency of HPV 67 in our results has not been investigated in larger meta-analyses.

In the meta-analysis by Guan et al. [27], African women with LSIL/ASCUS were HPV 58 positive in important proportions, and women with HSIL/ICC were predominantly infected by HPV 16 & 18, which is in agreement with our findings.

Our data from rural area cannot be disaggregated by cytological results but results are roughly similar to results found in Rwanda and Kenya [27;35]. Stratified by HIV status, the most frequent HPV types in our results are overall the same as results from a study done in Tanzania [32]. Our results highlight a high prevalence of the much known carcinogenic HPV types (HPV 16, 18, 52, 51, 56, 33 and 58) and very likely to persist in this unscreened population, which translates into high rates of cervical precursor lesions, making this population a high priority for public health interventions, regardless of their HIV-status. This is reinforced by the fact that this population has never been screened for cervical cancer lesions and clearly shows the need for implementation of cervical cancer screening intervention in Burundian women population.

In rural, the results in figure 2 imply that the Cervarix^®^ or Gardasil^®^ vaccines would target 32.1% and 30.8% of the HR/pHR-HPV infections among HIV-positive and negative women respectively. The nonavalent vaccine, Gardasil-9^®^, would target 60.7% and 53.9% of the circulating HR/pHR-HPV infections among HIV-positive and negative women respectively. In urban women, Gardasil^®^ or Cervarix^®^ vaccine would target 8.7% and 15.8% of all HR/pHR-HPV infections in HIV-positive and negative women respectively and Gardasil-9^®^ would prevent 31.9% and 42.1% of all HR/pHR-HPV infections in HIV-positive and negative women respectively. Overall, our results showed that Gardasil-9 vaccine covers most HR/pHR-HPV infections. If we look at the urban results, cervarix^®^ would target 2/3^rd^ and all HSIL/ASC-H/ICC among HIV-positives and negatives respectively; whereas Gardasil-9^®^ would target 7 on 8 and 100% of all HSIL/ASC-H/ICC among HIV-positives and negatives respectively (Table 2).

We are aware, however, of the described cross-protection [45] of HPV vaccines and studies on HPV strain distributions in invasive cervical cancer cases are also needed. The results of these studies plus the related cost-effectiveness studies are necessary elements to provide Burundian health authorities a sound evidence to choose the appropriate HPV screening and/or vaccination programmes.

### Study limitations

Our study population choose 4 similar strata according to HIV status and urban/rural and participants were not randomly selected. Thus, our overall study results do not represent the general female population of Burundi. This was a deliberate choice as we wanted to avoid that urban women and HIV infected women would be underrepresented. According to the 2010 Burundi Demographic Health Survey, 91% of Burundian women live in rural areas where HIV prevalence is estimated at 1.2% in women aged 15-49 years old, we assume that the results of the HIV negative women in Kirundo may be most representative result for the Burundian situation. This limitation turns indeed out to be a strength as we provide comprehensive baseline estimates of HPV prevalence and type-specific distribution in HIV-positive and negative women in rural and urban Burundi. And we observed the limited additional risk of rural HIV-infected women (who are relatively few) and the increased risk of the urban HIV-positive women. Another limitation is related to the use of two different HPV tests for rural and urban data which might have different sensitivity. Also, cytology results were only available for urban population with relatively few numbers across different cytological strata. Thus, caution is required for the inference of our results to the general population.

### Conclusions

In Burundi there is a high burden of HPV infection, in particular HR/pHR-HPV infections which mimics other similar settings. We observe that HIV-infected women in Bujumbura are a specific risk group to be targeted in contrast to the rural area where all women seem to have similar distributions of HPV circulating strains. This study strengthens the urgent need to introduce a comprehensive cervical cancer control program adapted to the context. This most probably may include a HPV-vaccination programme though cannot be the sole component due to the actual burden, the coverage and efficacy and lag time of the impact. Complementary studies in invasive cervical cancer cases are needed to further elucidate the most appropriate control tools.

## Acknowledgements

We would like to sincerely thank all study participants for their cooperation. We also thank Dr Christian Ngabirano, Dr Franck Kavabushi, CPAMP and ABUBEF teams for their support during data collection and support to get in contact with study participants in this study. Last but not least, we also acknowledge the VLIR-IUC/UB and VLIR-WAKA for their financial support in this study.

## List of abbreviations

ABUBEF: Association Burundaise pour le Bien-Etre Familial
AGC: Atypical Glandular Cells
ANSS: Association Nationale de Soutien aux Séropositifs et malades du SIDA
ASC-H: Atypical Squamous Cells cannot exclude High-grade lesion
ASCUS: Atypical Squamous Cells of Undetermined Significance
CI: Confidence Interval
CIN: Cervical Intraepithelial Neoplasia
DNA: DesoxyriboNucleic Acid
HIV: Human Immunodeficiency Virus
HPV: Human Papillomavirus
HR/pHR-HPV: High-risk/possible high-risk human papillomavirus
HSIL: High-grade Squamous Intraepithelial Lesions
ICC: Invasive Cervical cancer
LMICs: Low and Middle Income Countries
LR-HPV: Low-risk human papillomavirus
LSIL: Low-grade Squamous Intraepithelial Lesions
OR: Odds ratio
PCR: Polymerase Chain Reaction
S.D: Standard Deviation
SSA: Sub-Saharan Africa
STI: Sexually Transmitted Infection
VIA: Visual Inspection with 5% Acetic acid
VILI: Visual Inspection with Lugol’s Iodine
WHO: World Health Organization

## Declarations

### Availability of data and material

The dataset used and/or analysed during the current study are available from the corresponding author on reasonable request.

### Authors’ contributions

Conceptualisation: ZN, JPVG, DVB

Data curation: ZN, JPV

Analysis and interpretation of data: ZN, JPVG, DVB

Investigation (laboratory part): DVB, LRL, JB, IB

Methodology: ZN, JPV, DVB

Wrote or reviewed and approved the final manuscript: ZN, JPVG, DVB, LRL, JB, IB

